# Carbon and nitrogen availability affect biofilm growth and morphology of the extremotolerant fungus *Knufia petricola*

**DOI:** 10.64898/2026.03.19.712823

**Authors:** Abolfazl Dehkohneh, Julia Schumacher, Bastiaan J. R. Cockx, Karin Keil, Tessa Camenzind, Jan-Ulrich Kreft, Anna A. Gorbushina, Ruben Gerrits

## Abstract

Rock-inhabiting fungi thrive in subaerial oligotrophic environments such as desert rocks, solar panels and marble monuments where organic carbon and nitrogen are scarce. We tested whether the rock-inhabiting fungus *Knufia petricola* showed a preference regarding nitrogen (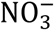 or 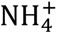) and carbon (glucose or sucrose) sources and whether it was sensitive towards carbon and nitrogen limitation. As this fungus produces the carbon-rich, nitrogen-free 1,8-dihydroxynaphthalene (DHN) melanin, we tested whether a melanin-deficient mutant would be less sensitive to carbon limitation. The carbon and nitrogen concentrations were the primary predictors of growth, with a broad optimum partially explained by an optimal fungal C:N ratio. Limiting carbon or nitrogen supply decreased biomass formation, CO_2_ production and biofilm thickness but promoted substratum penetration through filamentous growth. The nitrogen content of the biomass was flexible within limits, increasing upon increasing nitrogen supply or decreasing carbon supply. The carbon use efficiency was fairly constant, whereas melanization correlated with a higher nitrogen content of the biomass despite melanin being nitrogen-free. In conclusion, *in vitro*, *K. petricola* switches to explorative growth under nutrient limitations, like fast-growing fungi, revealing universal fungal resource-acquisition patterns.

**Graphical abstract text and image:** Carbon and nitrogen availability affect biofilm growth and morphology of the extremotolerant fungus *Knufia petricola*

Abolfazl Dehkohneh, Julia Schumacher, Bastiaan J. R. Cockx, Karin Keil, Tessa Camenzind, Jan-Ulrich Kreft, Anna A. Gorbushina, Ruben Gerrits

Growth of the rock-inhabiting fungus *Knufia petricola* was studied by varying carbon and nitrogen sources and concentrations. Overall, growth was best predicted by the carbon and nitrogen concentrations. Carbon and nitrogen limitation promoted substratum penetration through filamentous growth.

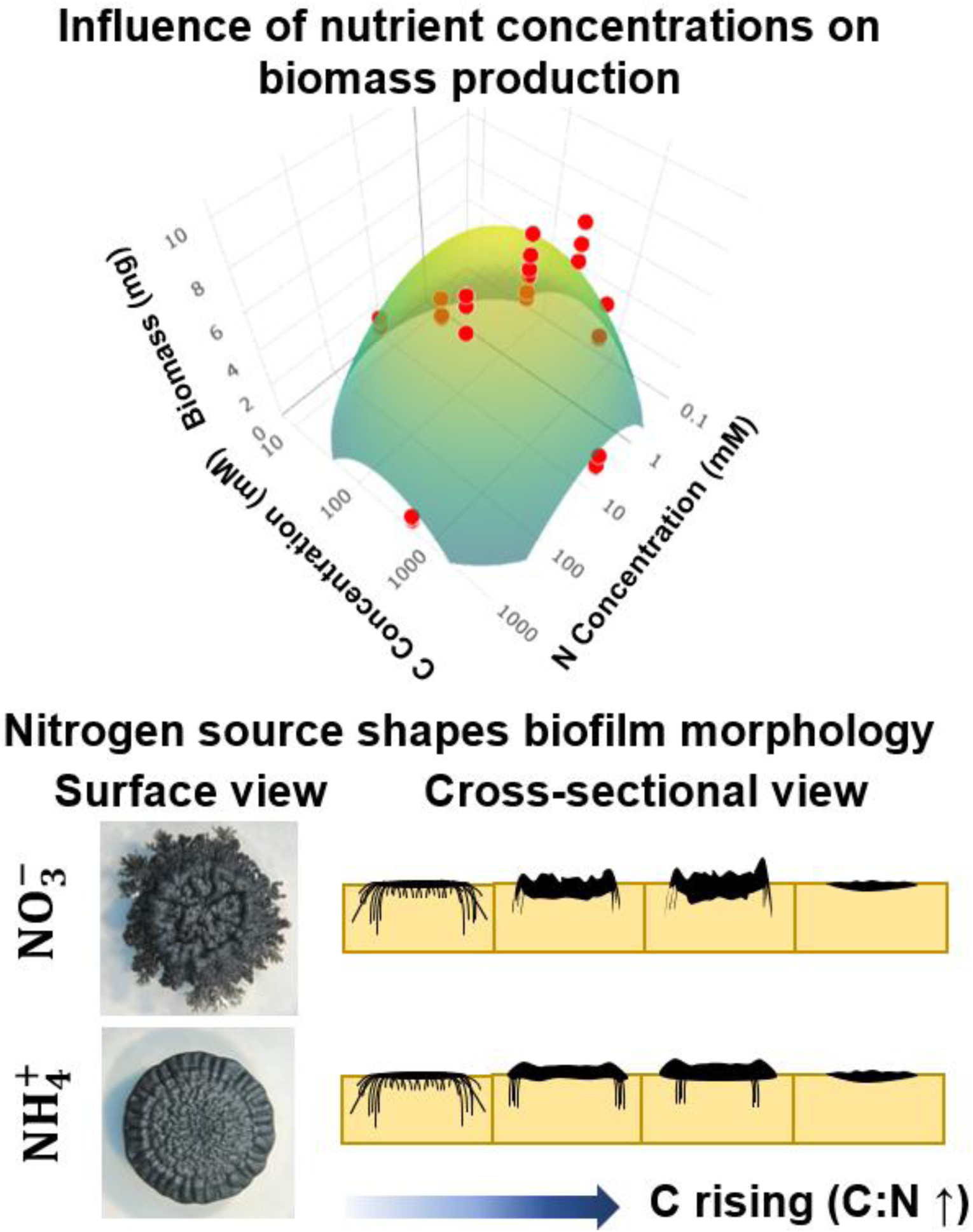

## 1. Introduction

Oligotrophic habitats that are exposed to air (i.e., subaerial habitats) are characterized by chronically low concentrations of bioavailable nutrients, particularly carbon and nitrogen (Gadd, 2007; Gorbushina, 2007). In such settings, organic carbon can originate from multiple sources, including dust−typically richer in organic carbon than the parent soil from which the dust originated (Webb et al., 2013)−as well as aerosols (Vodička et al., 2025), rain (Willey et al., 2000), volatile organic compounds (Guenther, 1997), pollen (Vogel et al., 2024) or phototrophic symbionts. Nitrogen can be delivered through rain and aerosols (Cornell et al., 2001), gaseous ammonia in agricultural settings (Robarge et al., 2002), gaseous nitrate in urban settings (Lin et al., 2006), or nitrogen-fixing cyanobacteria (Fujita and Uesaka, 2022). Absolute amounts of both carbon and nitrogen remain low because these surfaces have minimal *in situ* primary production, episodic and small external inputs, low sorption/retention of organics and ammonium on mineral phases, making microbial demand frequently exceed supply (Gadd, 2007; Gorbushina, 2007; Dang and Chen, 2017; Chu et al., 2025). Fungi colonizing these nutrient-poor environments are important contributors to nutrient cycling, decomposition of organic substrates, and the regulation of biogeochemical cycles. They deploy specialized strategies to do so, such as morphological and physiological adaptations (Gadd, 2007; Gorbushina, 2007; Liu et al., 2022).

One group of fungi able to colonize oligotrophic, subaerial surfaces are extremotolerant rock-inhabiting fungi (RIF), which can be isolated from surfaces like roof tiles (Ruibal et al., 2018; Prenafeta-Boldú et al., 2022) and marble monuments (Gorbushina et al., 2008). RIF are adapted to survive the multiple stresses typical for these environments such as nutrient scarcity (Gorbushina, 2007; Gostinčar et al., 2012), and osmotic, radiation and oxidative stresses (Aslanidi et al., 2003; Kogej et al., 2007; Fernandez and Koide, 2013; Gostinčar and Gunde-Cimerman, 2024). These fungi typically display three cell morphologies: yeast-like, meristematic, and (pseudo)hyphal (Gorbushina et al., 1993; Wollenzien et al., 1995; Chertov et al., 2004), between which they may switch depending on environmental conditions (Tonon et al., 2021). RIF are furthermore characterised by their generally slow growth and constitutive production of the black pigment melanin (Gorbushina et al., 1993; Gorbushina, 2007; Kogej et al., 2007). Melanin, being mostly incorporated into the outer cell wall, has been shown to enable their resistance to several stresses (Fernandez and Koide, 2013; Casadevall et al., 2017; Catanzaro et al., 2024a).

Most fungi can assimilate a wide range of nitrogen and carbon sources. Regarding carbon, simple carbohydrates, particularly glucose, are more easily utilized than more complex sugars such as sucrose (Ruijter and Visser, 1997). Glucose supports faster growth as its uptake is more rapid and its entry into central metabolic pathways is more direct (Nehls, 2008; Mäkelä et al., 2018). Regarding nitrogen, many species prioritize ammonium over nitrate (Celar, 2003), as the reduction of nitrate to ammonium requires eight electrons (reducing ATP production in the respiratory chain) as well as the synthesis of additional enzymes (Campbell and Kinghorn, 1990; Tudzynski, 2014).

The carbon-to-nitrogen (C:N) ratio of the medium is a primary determinant of the fungal growth. Individual species exhibit an optimal C:N ratio for their biomass production (Camenzind et al., 2020; Han et al., 2024). Increasing the C:N ratio by reducing mineral nitrogen supply creates nitrogen scarcity that redirects resources toward nitrogen acquisition. This shift has been shown to increase mycelial extension (Camenzind et al., 2020), reduce mycelial density (Camenzind et al., 2020), and lower overall biomass yield (Di Lonardo et al., 2020). These *in vitro* findings contrast with field studies, where fungal abundances generally decrease with increasing nitrogen additions to soils (de Vries et al., 2006; Wallenstein et al., 2006; Riggs and Hobbie, 2016). Such apparently negative responses of soil microbes to nitrogen additions are potentially driven by indirect effects, i.e., shifts in pH or soil-plant interactions (Treseder, 2008), making the interpretation of microbial nutrient limitation studies challenging.

For extremotolerant black fungi, knowledge regarding how different carbon and nitrogen sources and their relative proportions affect growth performance and biofilm morphology remains limited. Moreover, melanin, a defining trait of these fungi, may play roles beyond stress resistance, potentially influencing physiological adaptation to nutrient conditions.

Here, we aimed to address these gaps, by focusing on *K. petricola*, a model for RIF with a sequenced genome and available genetic engineering tools allowing functional genetics studies (Voigt et al., 2020; Erdmann et al., 2022; Erdmann et al., 2024; Schumacher, 2024). We evaluated (1) how carbon (glucose vs. sucrose) and nitrogen (nitrate vs. ammonium) sources and concentrations (and thus varying C:N ratios) affected growth parameters such as penetration depth, biomass production and carbon use efficiency (CUE), and (2) whether the formation of carbon-rich, nitrogen-poor melanin affected these nutrient-dependent growth characteristics by comparing the melanized wild type (WT) with a non-melanized mutant (Δ*pks1*). We hypothesized that *K. petricola* as a extremotolerant fungus would be able to grow at low carbon and nitrogen supplies, would have no preferences for the types of carbon or nitrogen sources and, if melanized, would be more sensitive to carbon limitation. In short, we found that carbon and nitrogen limitation affected the growth characteristics of *K. petricola* most strongly, while melanin formation and the type of carbon or nitrogen source were less important.

## 2. Materials and methods

### 2.1. General experimental design

The effects of glucose (Glc) and sucrose (Suc) as carbon sources were tested in combination with NaNO_3_(hereafter referred to as 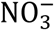) and NH_4_Cl (hereafter referred to as 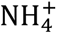) as nitrogen sources. The molar C:N ratios were adjusted by manipulating either the carbon or the nitrogen concentration of a defined minimal medium; these conditions are hereafter referred to as the carbon and nitrogen experiment. In the carbon experiment, the 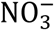 and 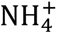 concentrations were kept at 10 mM while the Glc or Suc concentrations were varied (Table 1). In the nitrogen experiment, the carbon concentration was kept constant, at 100 mM for Glc or 50 mM for Suc, while the nitrogen supply was varied (Table 1). The medium was supplemented with the following salts and trace elements, at final concentrations of 5.88 mM KH_2_PO_4_, 6.71 mM KCl, 4.15 mM MgSO_4_, 0.90 mM CaCl_2_, 0.16 µM H_3_BO_3_, 0.025 µM MnCl_2_, 7.55 µM FeSO_4_, 0.80 µM CoCl_2_, 0.10 µM NiCl_2_, 0.059 µM CuCl_2_, 0.50 µM ZnSO_4_, 10 µM NaOH, 0.023 µM Na_2_SeO_3_, and 0.149 µM Na_2_MoO_4_, diluted in ultrapure water (Milli-Q). Five control media were prepared in which the carbon source, the nitrogen source, or both the carbon and nitrogen sources were omitted (Table 1). The pH was adjusted to 5. Media were solidified by adding 15 g L^-1^ bacteriological agar (AppliChem) and sterilized via autoclaving.

**Table 1.** Concentration of nutrients in the carbon and nitrogen experiments. In the carbon experiment, the concentration of the nitrogen source, either 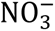 or 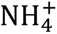, was constant at 10 mM and the concentration of carbon was varied from 1 mM to 1 M for Glc and 0.5 mM to 500 mM for Suc. In the nitrogen experiment, the carbon concentration was constant, either 100 mM for Glc or 50 mM for Suc, while nitrogen concentration was varied from 0.1 mM to 1 M.

| C:N ratio (molar) |  | 0.6 | 6 | 60 | 600 | 6000 | Controls |  |  |
| --- | --- | --- | --- | --- | --- | --- | --- | --- | --- |
| Carbon experiment | | | | | | | C -/ $\text{NO}_3^-$ | C -/ $\text{NH}_4^+$ | C -/ N - |
| C | Glc (mM) | 1 | 10 | 100 | 1000 |  | 0 | 0 | 0 |
|  | Suc (mM) | 0.5 | 5 | 50 | 500 |  | 0 | 0 | 0 |
| N | $\text{NO}_3^-$ (mM) | 10 | 10 | 10 | 10 | | 10 | | 0 |
| | $\text{NH}_4^+$ (mM) | 10 | 10 | 10 | 10 | | | 10 | 0 |
| Nitrogen experiment |  |  |  |  |  |  | Glc/ N - | Suc/ N - |  |
| C | Glc (mM) | 100 | 100 | 100 | 100 | 100 | 100 |  |  |
|  | Suc (mM) | 50 | 50 | 50 | 50 | 50 |  | 50 |  |
| N | $\text{NO}_3^-$ (mM) | 1000 | 100 | 10 | 1 | 0.1 | 0 | 0 | |
| | $\text{NH}_4^+$ (mM) | 1000 | 100 | 10 | 1 | 0.1 | 0 | 0 | |

### 2.2. Strains and culturing

Experiments were performed with *K. petricola* strain A95, isolated from a marble monument in Athens, Greece (Gorbushina et al., 2008). The melanin-deficient mutant Δ*pks1* (KP-0033, transformant PN3) was described previously (Voigt et al., 2020). Prior to experiments, the strains were inoculated onto solid minimal medium containing 100 mM Glc and 10 mM 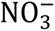 (i.e., a C:N ratio of 60) and incubated for 7 days at 25℃ in darkness. The fungal biofilms were collected and, together with 500 µL ultrapure water and 4-5 glass beads (3 mm diameter), mechanically dispersed for 5 min using a mixer mill (Retsch) set at 30 s^-1^. Colony forming units (CFU) in the suspensions were determined using a Thoma cell counting chamber and adjusted to a final titer of 10⁷ CFU mL^-1^. Ten µL of these inocula were separately dropped onto 12.5 mL of solid medium in 6-cm Petri dishes and incubated for 28 days at 25℃ in darkness.

### 2.3. Biofilm morphology

On day 28, micrographs of the biofilms and their cross-sections were taken using a digital camera and a stereomicroscope (Stemi 2000-C, Carl Zeiss). Cross-sections (∼2 mm thick) were cut out from the middle of the biofilms using a scalpel and placed on microscope slides for imaging. Cross-sections images were used to measure the maximal thickness of the biofilm and the maximal depth of visible penetration into the agar. For this purpose, the agar surface next to the biofilm was used as a reference height relative to which thickness and penetration were quantified. When biofilm growth pushed the agar upwards, the upper surface of this agar was used as the reference height.

The biofilm size was quantified using the Feret diameter, the longest distance between any two points on the biofilm edge, via ImageJ (Schneider et al., 2012). This was measured for three distinct zones of the biofilms, the innermost inoculation zone, the surrounding zone with compact growth and the outermost zone with filamentous growth (Figure 1). The inoculation zone is the extent of the biofilm measured after inoculation (day 0). The compact growth zone is dense and lacks visible filaments, and the filamentous growth zone is situated around the compact growth zone and characterised by the presence of filaments (Figure 1).

**Figure 1.**
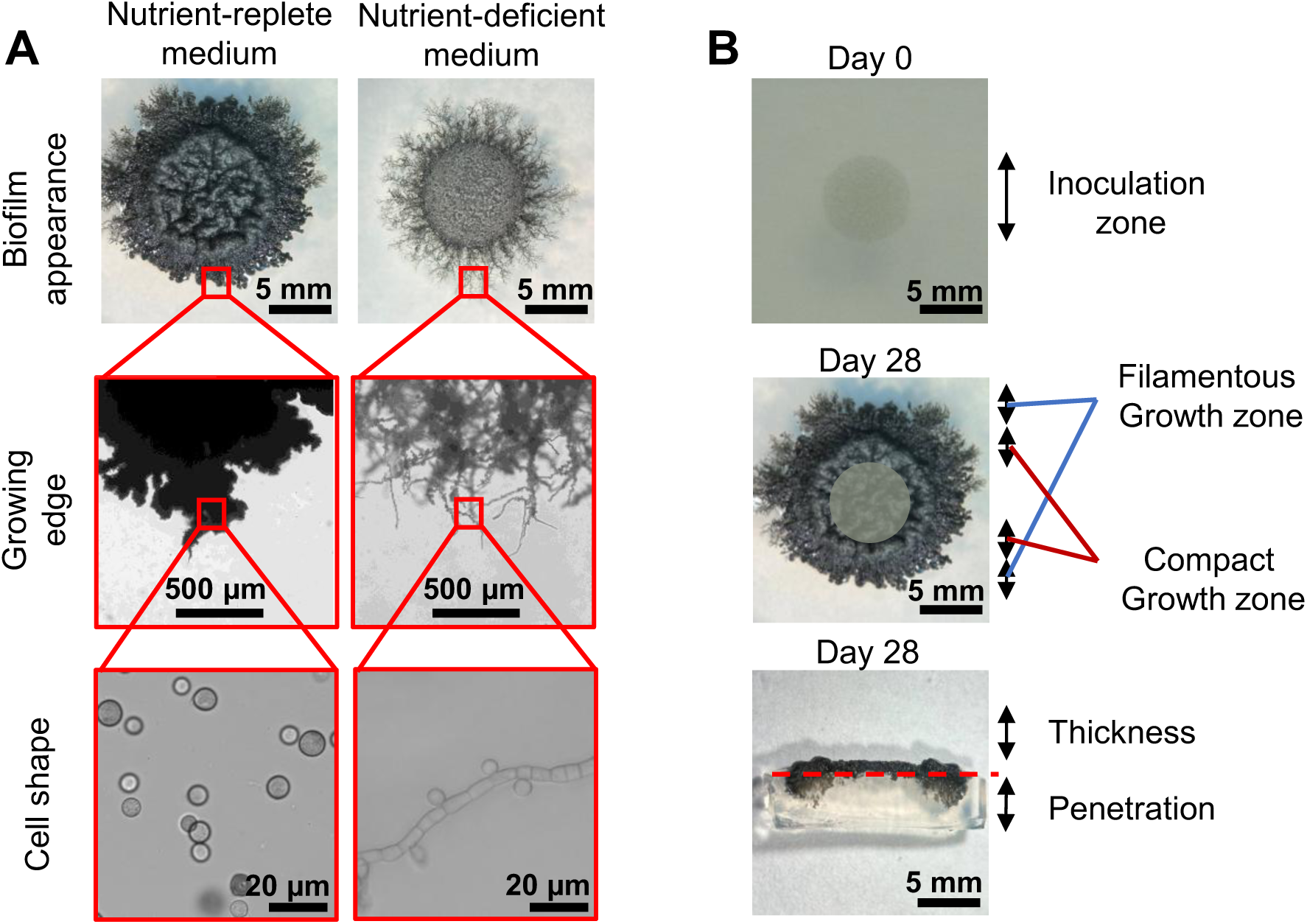
The morphology of *K. petricola* biofilms and cells depended on nutrient availability. Droplets containing 10^5^ CFU of *K. petricola* (WT) were spotted onto solid media with defined nutrient contents and incubated for 28 days at 25°C in the dark. **(A)** Biofilms formed in nutrient-replete (100 mM Glc and 10 mM 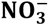) or nutrient-deficient (1 mM Glc and 0.1 mM 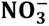) conditions. Starvation caused more filamentation around the biofilm edge. Yeast-like cells were observed in nutrient-replete conditions while more filaments were present in nutrient-deficient media. **(B)** Definition of growth zones. The maximum sizes of the inoculation zone and the compact and filamentous growth zones were measured to study the effects of nutrient availability on biofilm morphology. Penetration depth and biofilm thickness were quantified relative to the agar surface adjacent to the biofilm. Growth above this reference plane was recorded as biofilm thickness, whereas growth below was recorded as penetration depth.

### 2.4. Biofilm biomass

For determining the biomass on day 28, biofilms were scraped from the agar using a scalpel and transferred separately to bottles containing 50 mL of ultrapure water. The bottles were microwaved for 5-7 min to melt the agar. The resulting mixture was poured into 50-mL tubes and the biofilm was allowed to separate from the aqueous agar solution. Once the biofilm settled at the bottom (no centrifugation necessary), the supernatant (i.e., water and dissolved agar medium) was discarded, and the biofilm was washed twice with hot water to remove residual agar. Afterwards, the biofilms were washed, transferred to 2-mL polypropylene microcentrifuge tubes, and dried in a heating cabinet at 65℃ for two days. Subsequently, the tubes were equilibrated to room temperature and the dry biomass weighed. Partial loss of filaments could have occurred during the unavoidable heating and washing steps, but most of the biomass was concentrated in the denser zones and therefore unlikely to get lost.

### 2.5. Biofilm carbon and nitrogen contents

After weighing, the biomass samples were used for elemental analysis, pooling two technical replicates within each condition prior to measurement. Carbon and nitrogen contents were determined using an elemental analyzer (EA 3100, Eurovector) based on the Dumas combustion technique. Approximately 1-2 mg of biofilm sample was encapsulated in a tin capsule and subjected to dynamic flash combustion at 950℃ in a stream of pure oxygen. The resulting combustion gases (N_2_, CO_2_ and H_2_O) were passed through a reduction column and a moisture trap, followed by gas chromatographic separation and detection using a thermal conductivity detector. Elemental concentrations were quantified using the Weaver software supplied with the elemental analyzer (Eurovector).

### 2.6. Respiration and carbon use efficiency

Respiration was measured via regular CO_2_ concentration analysis of the headspace of sealed 100-mL glass vials containing 12.5 mL solid medium (as for the Petri dish experiments) and the fungal inoculant. The cultures were incubated for ten days in darkness at 25°C. The amount of CO_2_ gas produced was measured on day 1, 2, 3, 4, 7, 8, 9, and 10 using a LI-6400XT system (LI-COR). To avoid O_2_ limitation and CO_2_ accumulation, the vials were flushed for 5 min with ambient air that was rendered CO_2_-free by passing it through a soda-lime CO_2_ scrubber prior to entering the vials. Dissolved inorganic carbon was estimated from measured headspace CO_2_ concentrations using Henry’s law (Sander, 2015) at 25°C, treating dissolved CO_2_ and carbonic acid (H_2_CO_3_) as a single species. The partitioning between dissolved CO_2_ and bicarbonate (HCO^−^) was calculated using the apparent first dissociation constant of carbonic acid (pK_a1_ = 6.35) assuming that the pH was 4, as *K. petricola* can acidify the medium (Gerrits et al., 2020). Repeating the calculation for other pH values (i.e., 3 and 5) hardly changed the results. Total CO_2_ production was calculated as the sum of headspace CO_2_ and dissolved inorganic carbon. The carbon-use efficiency (CUE) was determined by dividing the carbon assimilated into biomass on day 10 (B) by the total carbon (R+B) that was either respired by, or assimilated into, the biomass according to the formula described by Maynard et al. (2017): *CUE* = *B*⁄(*R* + *B*). For some conditions, due to poor growth within 10 days, the CUE could not be calculated.

### 2.7. Statistical analysis

Most figures were created in Origin 2023. Asterisks in figures and tables indicate significance levels: * for p < 0.05, ** for p < 0.01 and *** for p < 0.001. All quantitative values are presented as mean ± standard error (SE). All statistical analyses and model visualizations were conducted in R (4.3.0) using the following R packages: readxl, ggplot2, plotly, corrplot, polycor, FactoMineR, GenSA, MASS, htmltools, dplyr, tidyverse and tidyr.

The dataset consisted of three biological replicates for each treatment combination in each experiment. The categorical variables strain, nitrogen source, and carbon source were treated as factors, while carbon and nitrogen concentrations or the C:N ratio were treated as quantitative variables. Quadratic terms for carbon and nitrogen concentrations or the C:N ratio were included to capture non-linear relationships expected from visual exploration of the data. To assess the influence of these explanatory variables on biofilm responses, distinct regression models were applied for each response variable. These initial, full regression models included all main effects and all possible interaction terms. Stepwise model reduction was performed in both forward and backward directions based on minimizing the Akaike Information Criterion (AIC) to identify the most parsimonious model. Model residuals were checked for normality, homoscedasticity, and independence to check the validity of assumptions.

We first compared models using either the C:N ratio or its log_10_-transformed values (including quadratic terms) as predictors, along with their interactions with strain, nitrogen and carbon treatment. These models were compared using statistical measures and physiological appropriateness. While for a few state variables the untransformed C:N ratio model achieved marginally higher adjusted R^2^, the shape of the fitted model indicated unphysiological predictions outside the data range and more extreme peaks and curvatures suggesting overfitting in the polynomial terms (10.5281/zenodo.18980984). Specifically, the model based on untransformed C:N ratios generated anomalous predictions (e.g., high peaks at C:N ratios ∼300 followed by steep declines). The log_10_-transformed C:N model produced more realistic broader and lower maxima rather than extreme behavior and therefore log-transformed data were used for further analysis.

Then, we compared two approaches before deciding on the most appropriate method. The first approach was to use the C:N ratio as an ‘integrative’ explanatory variable (called ratio-based models), and the other was to use carbon and nitrogen concentrations as separate explanatory variables (called concentration-based models). The former approach resulted in different regression models for both the carbon and nitrogen experiments, which made interpretation difficult as different optima were obtained for the carbon and nitrogen experiments. Moreover, although some response variables were heteroscedastic, the overall residual distributions were closer to the expected patterns for normal and homoscedastic variables for the concentration-based models. Note any systematic deviations from expected patterns were towards the ends of the range, which would not affect the regions around the optima. We therefore decided to use concentration-based models based on log_10_-transformed concentrations as the most appropriate models and these results are presented in the main text, for results from the other models see 10.5281/zenodo.18980984.

## 3. Results

### 3.1. Nutrient availability and type of nitrogen source controlled single cell and biofilm morphology

In general, the concentration of carbon and nitrogen affected both biofilm and single cell morphology of *K. petricola*. At higher concentrations of both nutrients, the biofilm was compact and consisted of yeast-like, spherical cells, while at lower concentrations, filamentous growth was observed surrounding the compact growth zone and consisted of pseudo-hyphal structures (Figure 1). The morphological analysis mostly indicated that the type of nitrogen source affected the shape of biofilms, regardless of the carbon source (Figure 2). In the presence of 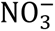, the biofilms were thicker at the edge, had small folds in the center and more filamentation, while in the presence of 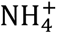, the biofilms had a smoother surface and edge and showed less filamentation (Figure 2A, B). However, at low concentrations of either carbon or nitrogen (C:N ratio of 0.6 in the carbon experiment and 6000 in the nitrogen experiment), the biofilm morphology was similar (Figure 2). At a C:N ratio of 60, penetration into the agar occurred for both 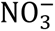 and 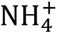. The biofilm morphologies of the WT and the Δ*pks1* mutant were similar. Not adding either carbon, nitrogen or both triggered filamentation around the compact growth zone (Figure 2C). Increasing the supply of nitrogen when carbon was limiting did not affect biofilm morphology, whereas increasing the carbon concentration when nitrogen was limiting resulted in better growth at intermediate Glc concentrations and dual nitrogen and carbon limitation at the lower Glc concentration of 1 mM (Figure S1).

**Figure 2.**
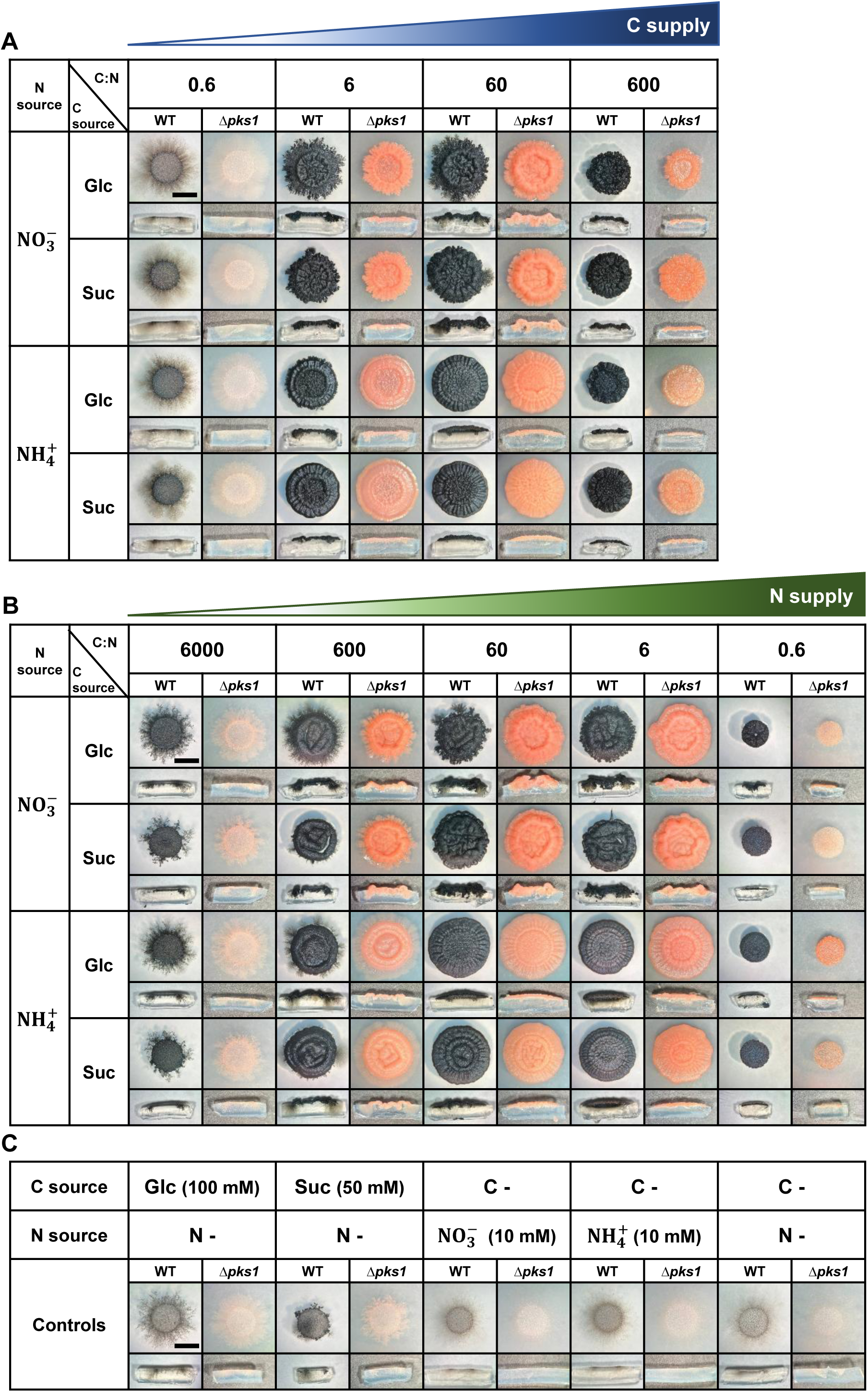
Nutrient levels and nitrogen source determine biofilm morphology of *K. petricola* strains. One droplet containing 10^5^ CFU of the WT or the melanin-deficient Δ*pks1* mutant was dropped on the solid medium and incubated for 28 days at 25°C in the dark. All experiments were conducted in triplicate; images of representative biofilms (top views and cross-sections) are shown. Scale bars represent 5 mm. **(A)** The effect of carbon availability (designated as the carbon experiment) was tested by increasing the glucose (Glc) concentration from 1 mM to 1 M or sucrose (Suc) from 0.5 mM to 500 mM, keeping 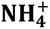 or 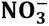 concentrations at 10 mM, corresponding to molar C:N ratios of 0.6 to 600. **(B)** The effect of nitrogen availability (designated as the nitrogen experiment) was tested by increasing the concentration of 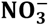 or 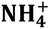 from 0.1 mM to 1 M, keeping the concentrations of Glc at 100 mM or Suc at 50 mM, corresponding to molar C:N ratios of 6000 to 0.6. **(C)** The effect of starvation was tested in unsupplemented media (C - /N -) or in media supplemented with a carbon or nitrogen source only.

The filamentous and compact growth zones increased and decreased, respectively, upon carbon or nitrogen limitation (Figure 3A, Figure 4A). The most parsimonious of the concentration-based regression models with log_10_-transformed carbon and nitrogen concentrations showed that the linear and quadratic terms of both carbon and nitrogen concentrations were strong predictors for both growth zones (Table 2 for main effects of the variables on growth responses). When carbon, nitrogen, or both were absent, biofilms expanded exclusively through filamentous growth, with no compact growth zone (Figure 3A and Figure 4A). The presence of 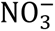 altered the nitrogen concentration–dependent response and promoted filamentous growth at intermediate nitrogen concentrations (Figure 3A, Figure 4A, Table S1 for interaction effects of the variables on growth responses), whereas 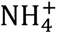 favored compact growth regardless of the nitrogen concentration (Table 2). Moreover, the WT produced more filaments than the mutant in presence of 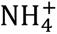 and Glc; however, the effect of strain depended on the nutrient source concentrations, as indicated by significant interaction terms in the model (Figure 3A, Figure 4A, Table 2, Table S1).

**Figure 3.**
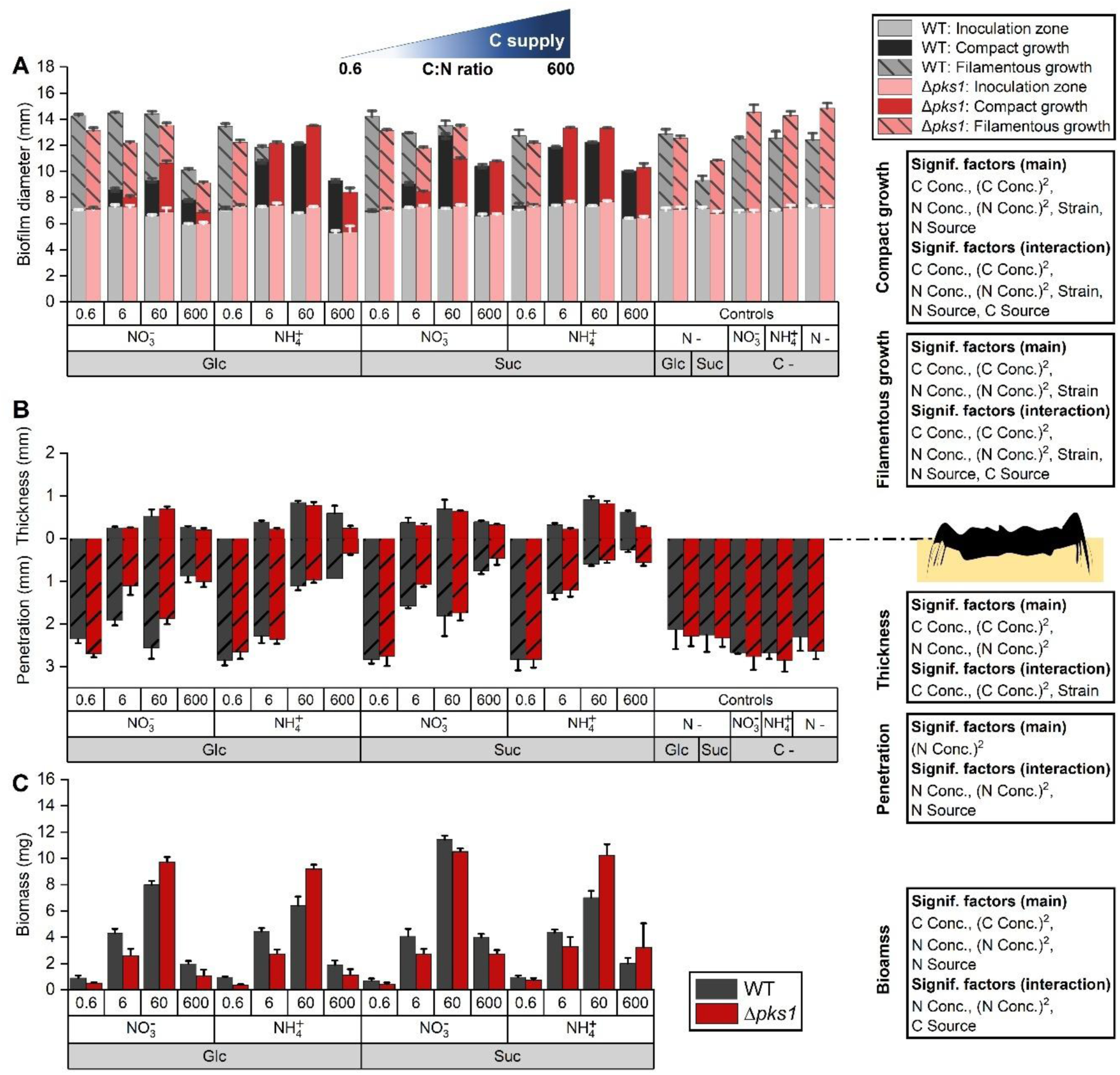
Effects of increasing carbon supply (the carbon experiment) on filamentation, substrate penetration, biofilm thickness and biomass for WT and *Δpks1* mutant after 28 days. **(A)** Increasing the carbon concentration reduced filamentation and increased compact growth (both based on maximum diameter of respective zones). Absence of either carbon or nitrogen led to biofilms expanding via filamentation. **(B)** Biofilms grown on media with lower concentrations of carbon were thinner and penetrated the agar more deeply. **(C)** Biomass was highest at the intermediate C:N ratio of 60. The amount of biomass in the controls was too low to be measurable. All experiments were conducted in triplicate; error bars show SE. Statistical significance is shown for the main variables and for the interactions among these variables based on the most parsimonious regression models (Table 2).

**Figure 4.**
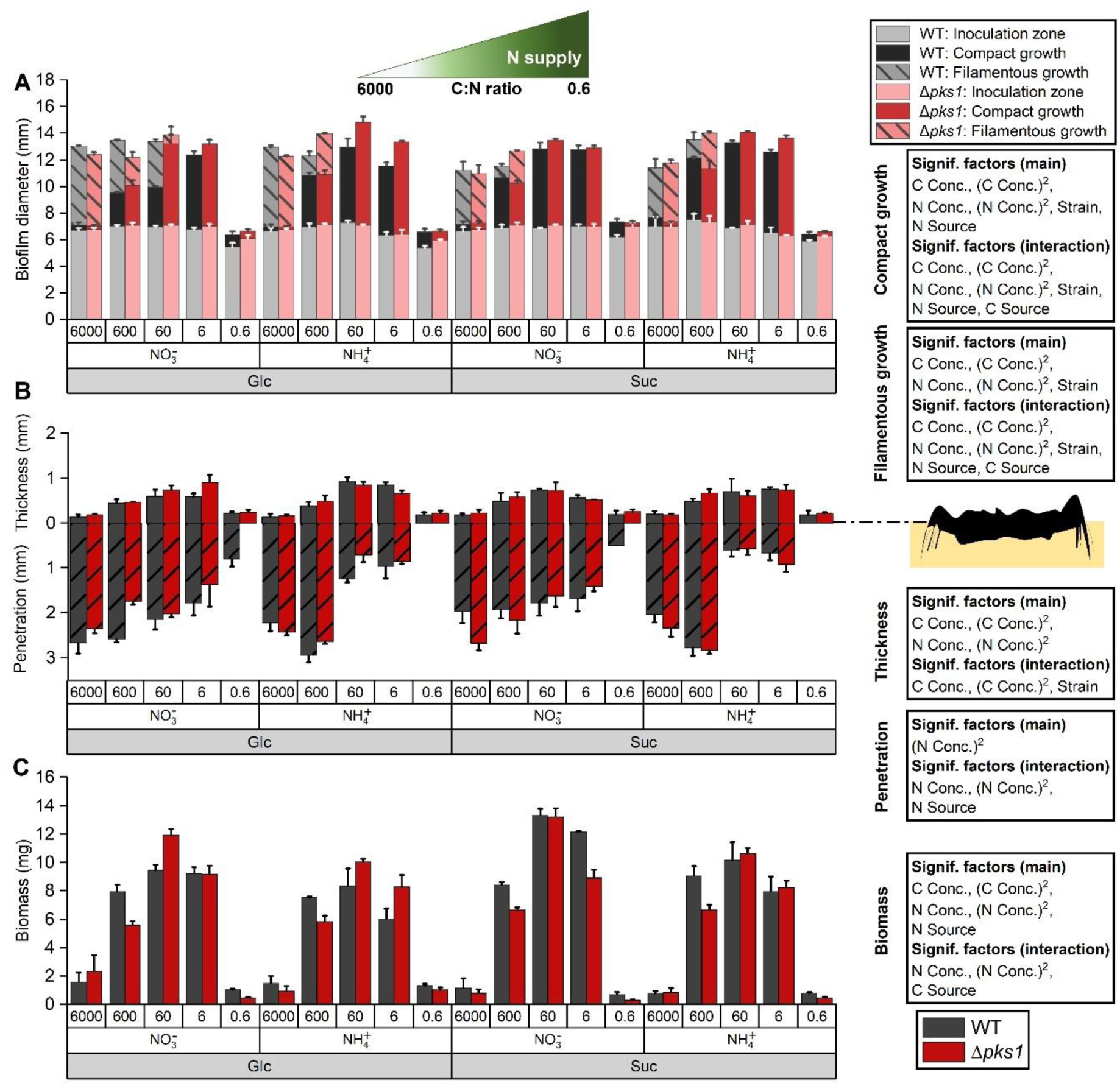
Effects of increasing nitrogen supply (the nitrogen experiment) on filamentation, substrate penetration, biofilm thickness and biomass for WT and *Δpks1* mutant after 28 days. **(A)** Increasing nitrogen concentration reduced filamentous growth and increased compact growth (both based on maximum diameter of respective zones). **(B)** Biofilms penetrated the agar more deeply at lower nitrogen concentrations while thickness increased with increasing nitrogen. **(C)** The highest biomass was formed at the intermediate C:N ratio of 60. All experiments were conducted in triplicate; error bars show SE. Statistical significance is shown for the main variables and for the interactions among these variables based on the most parsimonious regression models (Table 2).

**Table 2.** Summary of the most parsimonious concentration-based regression models for biofilm growth responses to explanatory variables. Regression models were fitted to assess the effects of log_10_-transformed atomic concentrations of carbon and nitrogen (linear and quadratic terms) and the factors strain, nitrogen and carbon source on growth. The log10-transformed nutrient concentrations were included as both linear and quadratic terms to allow for the apparent nonlinear responses. Model selection was performed using stepwise removal and addition of terms to minimize the Akaike Information Criterion (AIC), which identified the most parsimonious model that balanced explanatory power and model complexity. Shown with a grey background are the variables which were removed from this parsimonious model. The intercept represents the predicted response at zero carbon and nitrogen concentrations for the reference levels of strain, nitrogen and carbon sources, and therefore serves as a baseline. Model fit was also evaluated using the adjusted R^2^, which accounts for the number of predictors included. Statistical significance is indicated using standard thresholds (* p < 0.05, ** p < 0.01, *** p < 0.001). Data represent measurements at 28 or 10 dpi (days post inoculation). Overall, carbon and nitrogen concentrations and their quadratic terms were the most significant predictors of most growth characteristics. This table shows only the main effects, the full models with all interactions are included in Table S1. For model fitting and more details see the R notebook (10.5281/zenodo.18980984).

|  | Intercept | Log (C atoms) | (Log (C atoms))<sup>2</sup> | Log (N atoms) | (Log (N atoms))<sup>2</sup> | Strain | Nitrogen source | Carbon source | AIC | Adjusted R<sup>2</sup> |
| --- | --- | --- | --- | --- | --- | --- | --- | --- | --- | --- |
| Thickness | *** | *** | *** | *** | *** |  |  |  | -755 | 0.689 |
| Penetration | *** |  |  |  | *** |  |  |  | -261 | 0.719 |
| Compact growth zone | *** | *** | *** | *** | *** | * | * |  | 64 | 0.815 |
| Filamentous growth zone | *** | *** | *** | *** | *** | * |  |  | -75.7 | 0.893 |
| Biomass 28 dpi | *** | *** | *** | *** | *** |  | * |  | 336 | 0.723 |
| C content 28 dpi | *** |  |  |  | * |  | * |  | 677 | 0.469 |
| N content 28 dpi | *** | *** | *** |  | *** |  |  |  | 325 | 0.863 |
| H content 28 dpi | *** |  |  |  |  | * |  |  | 341 | 0.334 |
| Fungal C:N 28 dpi | * | *** | *** | * |  | * |  |  | 338 | 0.638 |
| CO<sub>2</sub> 10 dpi | *** | *** | *** | *** | *** |  | ** |  | -125 | 0.917 |
| CUE 10 dpi | *** |  |  | ** | ** |  |  |  | 262 | 0.359 |
| C content 10 dpi | *** | *** | ** | *** |  |  | ** | *** | 415 | 0.357 |
| Biomass 10 dpi | *** | *** | *** | *** | *** |  |  |  | -129 | 0.781 |

The penetration depth increased with a decrease of either carbon or nitrogen supply, while the thickness showed the opposite trend (Figure 3B, Figure 4B). The carbon and nitrogen concentrations and their quadratic terms affected the thickness, while the penetration depth was only affected by the quadratic term of the nitrogen concentration (Table 2). Although the main effect of the nitrogen source was not significant, its significant interaction with the nitrogen concentration shows that the influence of the nitrogen source on penetration depth was concentration-dependent (Table S1), with 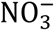 promoting deeper penetration under high nitrogen conditions (Figure 4B, Table 2, Table S1).

Overall, the carbon and nitrogen concentrations and their quadratic terms were the key predictors for biofilm morphology, with the nitrogen source having a weaker effect.

### 3.2. Nutrient availability determined biomass production

The biomass was affected by the carbon and nitrogen concentrations and their quadratic terms (Table 2), being highest at intermediate nutrient concentrations (Figure 3C, Figure 4C). For the nitrogen experiment, the biomass varied less within the C:N ratio range of 6 to 600 (Figure 4C) than for the carbon experiment.

Using the most parsimonious of the concentration-based regression models with log_10_-transformed carbon and nitrogen concentrations, the optimal biomass was inferred to be at 273 mM carbon and 10.1 mM nitrogen, i.e., a medium C:N ratio of 26.9, which is between the observed C:N ratios of 6 and 60 (Figure 5, Table S1).

**Figure 5.**
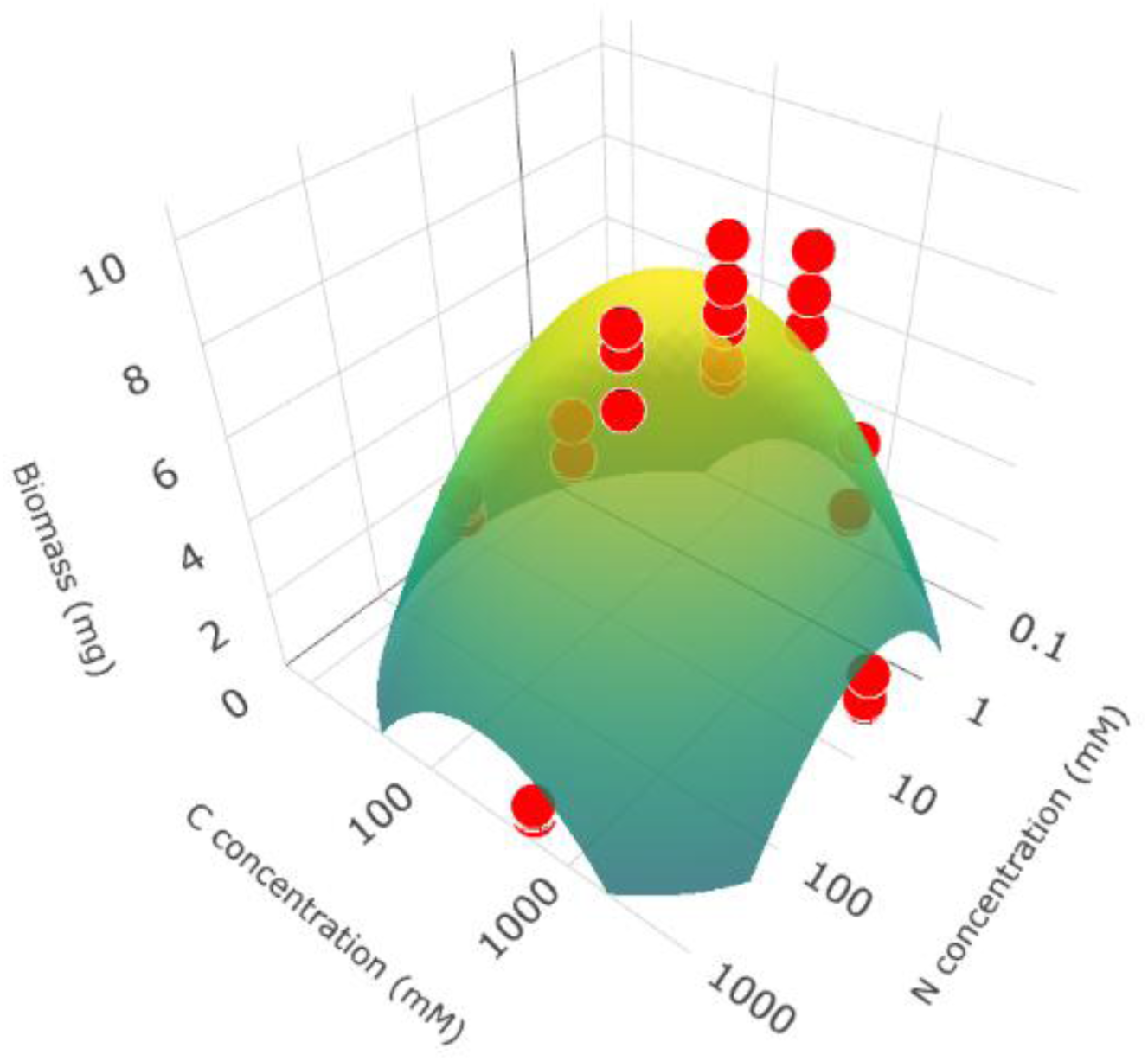
The model-inferred optimal C:N ratio of the medium for biomass growth. The surface visualizes the predictions from the most parsimonious (lowest AIC) concentration-based regression model using log₁₀-transformed atomic carbon and nitrogen concentrations as explanatory variables and biomass after 28 days as response variable. This model incorporated quadratic terms for both carbon and nitrogen concentrations to capture nonlinear dose-response relationships. In this concentration-based regression model, the carbon and nitrogen concentrations and their quadratic terms and the nitrogen source had a significant effect on the biomass (Table 2, Table S1). The surface shown is specific for the 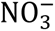 medium. Red dots show experimentally observed biomass values from both the carbon and nitrogen experiments for the WT grown on minimal medium with Glc and 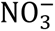 as the sole carbon and nitrogen sources. Axes display log₁₀-scaled carbon (C) and nitrogen (N) concentrations (mM), the z-axis indicates biomass (mg). The model predicts a broad biomass maximum at a C:N ratio of 26.9 (273 mM carbon and 10.1 mM nitrogen).

### 3.3. Biofilm carbon content was constant while nitrogen content was flexible

The carbon content of the biomass remained fairly constant upon changing the carbon and nitrogen concentrations, despite some small relative differences being significant (Figure 6A, C, Table 2). For instance, the WT biomass contained more carbon than the mutant at higher carbon concentrations (Figure 6C, Table S1).

**Figure 6.**
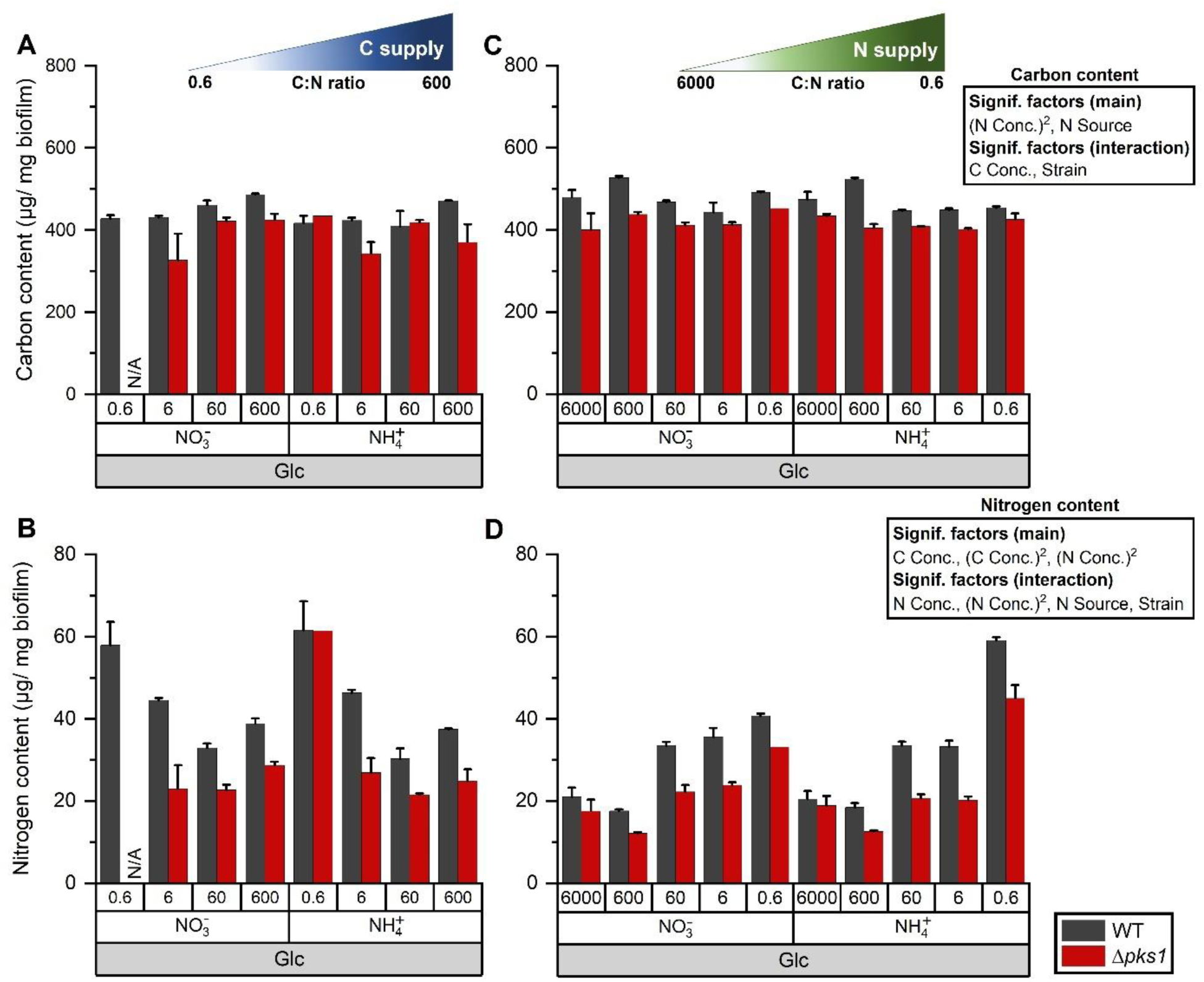
The carbon (A, C) and nitrogen (B, D) content of the fungal biomass of the WT and Δ*pks1* mutant in the carbon (A, B) and nitrogen (C, D) experiments. Varying both carbon and nitrogen concentrations of media had little effect on carbon content but affected the nitrogen content of biofilms, the latter decreasing upon increasing the carbon supply or decreasing the nitrogen supply. All experiments were conducted in triplicate, except the carbon experiment with the Δ*pks1* mutant at a C:N ratio of 0.6 with Glc and 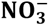 due to insufficient growth; error bars show SE. Statistical significance is shown for the main variables and for the interactions among these variables based on the most parsimonious regression models (Table 2).

The nitrogen content, however, was more strongly affected by the carbon concentration and its quadratic term, as well as by the quadratic term of the nitrogen concentration (Table 2), decreasing upon increasing the carbon supply or decreasing the nitrogen supply (Figure 6B, D). Moreover, upon increasing the nitrogen concentration, the amount of nitrogen in WT biofilms was higher than for the Δ*pks1* (Table S1).

The C:N ratio of the fungal biomass depended on the carbon concentration of the medium and its quadratic term, the nitrogen concentration and the strain (Figure S3, Table 2). With increasing carbon supply, the biomass C:N ratio increased to reach an observed optimum around the intermediate C:N ratio of 60. For lower nitrogen supply, the biomass C:N ratio was much higher than in the carbon experiment but dropped to ratios similar to the carbon experiment at higher nitrogen supply. Lastly, the biomass C:N ratio was higher for the melanin-deficient mutant than the WT.

Taken together, the WT biofilms had, for some experiments, higher amounts of carbon and nitrogen compared to non-melanized Δ*pks1* biofilms. The nitrogen content of the biofilm was strongly influenced by changing either carbon or nitrogen concentration, while the carbon content of the biofilm remained relatively stable.

### 3.4. Carbon use efficiency (CUE) was less affected by nutrient availability

Dedicated experiments were run for ten days on solid media in closed vials to determine the effect of carbon and nitrogen availability on respiration and CUE. Respiration, measured as cumulative CO_2_ production, generally peaked around intermediate carbon and nitrogen concentrations (Figure 7A and Figure 8A), being affected by the carbon and nitrogen concentrations and their quadratic terms (Table 2).

**Figure 7.**
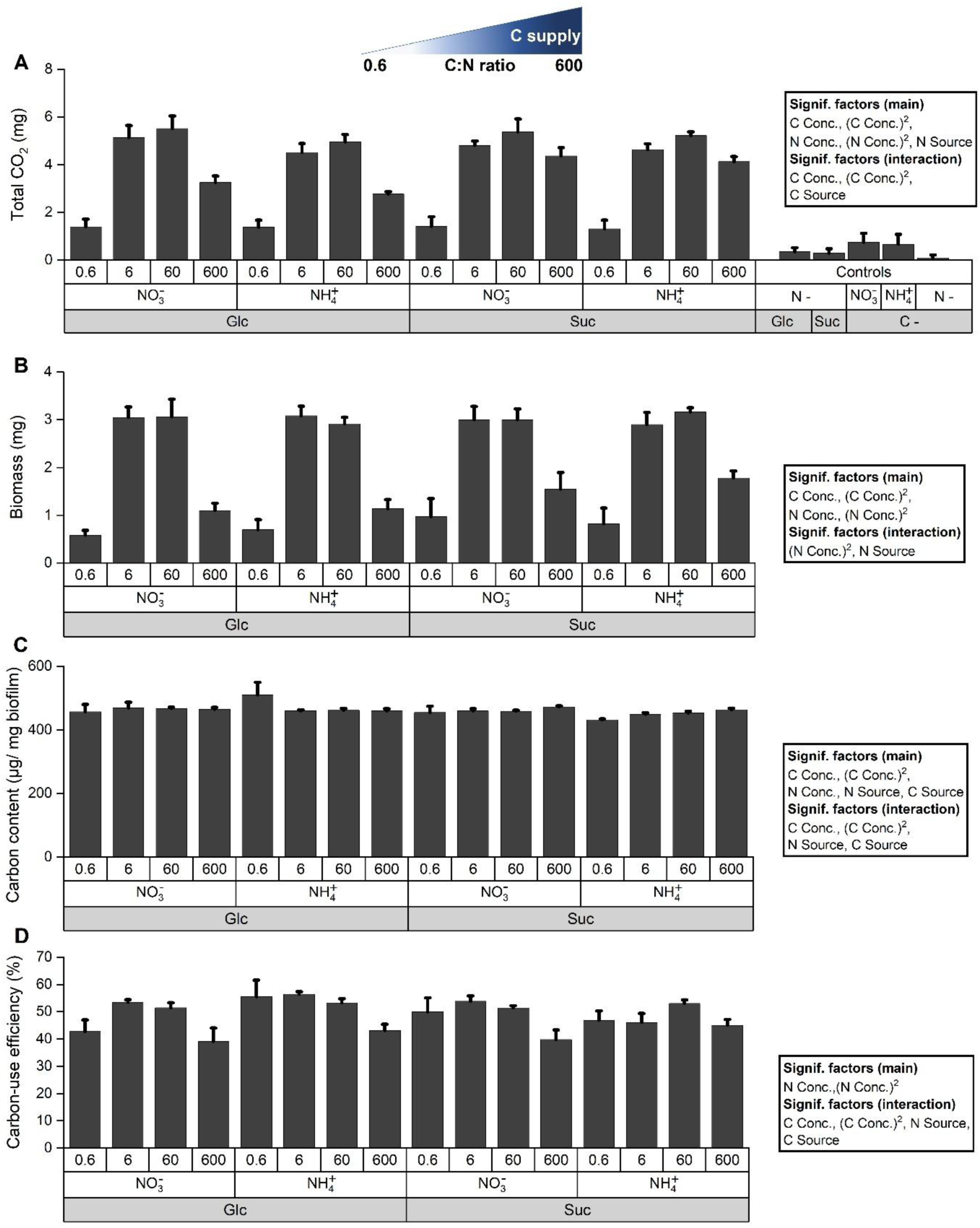
The effect of increasing carbon supply on respiration (**CO**_**2**_ production), biomass production, carbon content and carbon use efficiency (CUE). WT biofilms were grown in 100-mL vials containing 12.5 mL of solid medium at 25°C for 10 days. **(A)** Total CO_2_ (mg) produced over 10 days. Most CO_2_ was produced at intermediate carbon concentrations. Less CO_2_ was produced without nitrogen supplementation than without carbon supplementation. **(B)** Substantially more biomass was produced at intermediate carbon concentrations after 10 days. **(C)** Carbon content of the biomass after 10 days was only slightly increased at higher carbon concentrations. **(D)** CUE varied less than biomass and respiration. All experiments were conducted in triplicate; error bars show SE. Statistical significance is shown for the main variables and for the interactions among these variables based on the most parsimonious regression models (Table 2).

**Figure 8.**
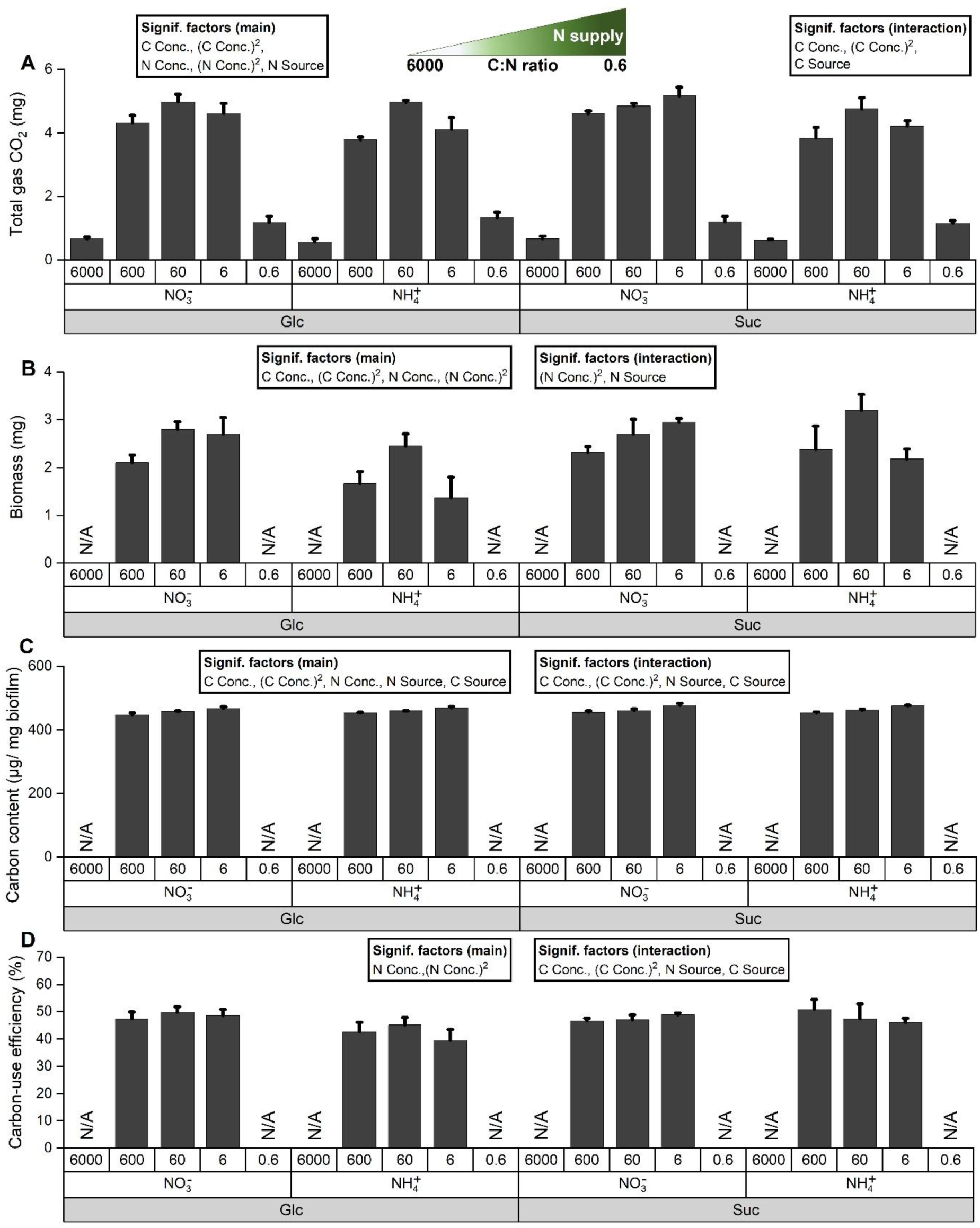
The effect of increasing nitrogen supply on respiration (CO_2_ production), biomass production, carbon content and carbon use efficiency (CUE). WT biofilms were grown in 100-mL vials containing 12.5 mL of solid medium at 25°C for 10 days. **(A)** Total **CO**_**2**_ (mg) produced over 10 days. More **CO**_**2**_ was produced at intermediate nitrogen concentrations. **(B)** Biomass production was also highest at intermediate nitrogen concentrations after 10 days. **(C)** The carbon content of the biomass increased slightly with the nitrogen concentration while **(D)** the CUE showed no clear trend. All experiments were conducted in triplicate; error bars show SE. Statistical significance is shown for the main variables and for the interactions among these variables based on the most parsimonious regression models (Table 2).

Biomass formation also peaked around intermediate carbon and nitrogen concentrations (Figure 7B, Figure 8B), likewise being affected by the carbon and nitrogen concentrations and their quadratic terms (Table 2).

Similar to the open Petri dish experiments, fungal carbon content in the closed-vial system showed only minor variation across carbon and nitrogen concentrations, although both linear and quadratic terms were statistically significant (Figure 7C and Figure 8C, Table 2).

The CUE was also quite similar for all conditions, being only marginally affected by the nitrogen concentration and its quadratic term (Table 2).

Overall, respiration and biomass formation had clear maxima around intermediate carbon and nitrogen concentrations while the carbon content and CUE were almost constant.

## 4. Discussion

Although numerous studies have investigated how carbon and nitrogen availability influence the growth of soil fungi, this work is, to our knowledge, the first to systematically dissect the effects of carbon and nitrogen sources and concentrations on a extremotolerant, rock-inhabiting fungus. Since these fungi are considered oligotrophic (Gostinčar et al., 2012), we hypothesized that they would be able to grow at low carbon and nitrogen supplies and would not have a preference for either type of carbon or nitrogen source. The use of the model rock-inhabiting fungus *K. petricola* enabled us to also study the effect of melanization by comparing the WT with the melanin-deficient Δ*pks1* mutant. We hypothesized that the production of this carbon-rich polymer would cause a higher sensitivity towards carbon limitation.

*K. petricola* grew to the highest observed extent at the intermediate medium C:N ratio of 60, as indicated by the largest compact growth zone, highest biomass production, thickest biofilm, and highest CO_2_ production. Other studies noted that the optimal observed medium C:N ratio varied considerably among fungal species: Hatakeyama and Ohmasa (2004) observed that Boletales species grew best at C:N ratios of 5 to 66, Gao et al. (2007) observed that Eurotiales and Hypocreales species grew best at C:N ratios of 10 to 40 and Di Lonardo et al. (2020) observed that a species of Mucorales and Hypocreales grew to the highest extent at a C:N ratio of 8. The reasons for these observed differences among species could include differences in species-specific plasticity of biomass C:N ratios, reduced CUE due to fermentation rather than respiration, use of different carbon and nitrogen sources or inaccuracies arising from unavoidable limitations in the number of tested C:N ratios. Camenzind et al. (2020) found that the model-inferred optima for 16 fungal species spanning multiple phyla, including Mucoromycota, Basidiomycota, and Ascomycota, were at medium C:N ratios ranging from 5 to 100, with an average at a C:N ratio of 47. These authors moreover noted that this ratio corresponds to twice the C:N ratio of the biomass of known fungal species and proposed that this is due to half the carbon being respired (CUE of 50%).

The same reasoning can be applied to our *K. petricola* data. For this fungus, the model-inferred optimal biomass was at the medium carbon concentration of 273 mM and nitrogen concentration of 10.1 mM, i.e., at a C:N ratio of 26.9 (based on all data from the carbon and nitrogen experiments for the WT growing on Glc and averaged for both nitrogen sources, Figure 5 shows results for 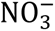). The model-inferred fungal C:N ratio (at this medium C:N ratio of 26.9) was 17.7 ± 2.4 (averaged over both nitrogen sources and the carbon and nitrogen experiments ± the 95% CI). Dividing this by the model-inferred CUE of 0.489 ± 0.031 (at the medium C:N of 26.9 and averaged over the same conditions), gives an estimate of the expected optimal medium C:N ratio of 36.2 ± 5.4 (Table S1). Since growth was not assessed at a medium C:N of 36.2, we need to compare the fungal C:N ratio and CUE-based expected optimum medium C:N ratio of 36.2 with the model-inferred optimal C:N ratio of 26.9. Since this optimum is broad rather than a sharp peak and was inferred from noisy data, it should not be considered to be exactly 26.9 and is thus broadly in line with the expected value of 36.2 ± 5.4. The lower-than-expected model-inferred biomass optimum could however indicate that our CUE was underestimated (expected and model-inferred figures would agree at a CUE of 0.66). Underestimating CUE could be due to loss of carbon through removal of carbon-rich extracellular polymeric substances (EPS) from the colonies during sample processing or due to CUE increasing over time (we only measured CUE at 10 days). Overall, this suggests that the optimal medium C:N ratio can be largely explained in terms of optimal fungal C:N ratio and CUE.

A variety of other ‘optima’ could be observed in the data (Table S1), but these maxima were more variable and dependent on the nitrogen source and strain. This is likely due to natural selection primarily optimizing biomass growth rather than the partially dependent variables such as thickness and compact growth zone.

While the compact growth zone, thickness, biomass and CO_2_ production decreased upon decreasing the supply of either carbon or nitrogen, thus following the Law of the Minimum (Sprengel, 1828), the filamentous growth zone and penetration depth increased upon lowering the supply of either nutrient. Filamentous growth was most extensive when both carbon and nitrogen were lowest. From the curvature of the fitted relationship (10.5281/zenodo.18980984), filamentous growth would likely increase further at even lower concentrations while it was absent around the optimal medium C:N ratio. These results suggest a nutrient scavenging role of filaments, which is also expected to be under direct selection. These observations disprove our hypothesis that this fungus would be less sensitive towards carbon and nitrogen limitations: *in vitro* (in the absence of physicochemical stresses) *K. petricola* behaves like fast-growing fungi. Interestingly, the total colony diameter did not change much, indicating that *K. petricola* prioritized extension for foraging over density under nutrient scarcity. Enhanced filamentation under nitrogen limitation is known to be common for fungi (Wallander and Nylund, 1992; Camenzind et al., 2020), but we are not aware of other studies showing enhanced penetration upon nitrogen limitation. Increased filamentation and penetration, however, clearly show that fungi have adapted to scavenge for limiting carbon or nitrogen. The decreased biomass and CO_2_ production under nitrogen limitation fits observations of other fungi (Camenzind et al., 2020; Di Lonardo et al., 2020). As filamentous growth and penetration would, based on the curvature of the fitted relationship (10.5281/zenodo.18980984), likely increase further at even lower concentrations, the ‘optima’ for this response are not given in Table S1.

Increasing the carbon and nitrogen supply beyond the C:N ratio of 60 (i.e., a C:N ratio of 600 for the carbon experiment and C:N ratios of 6 and 0.6 for the nitrogen experiment) by adding 1 M of Glc, 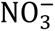 or 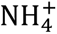 lowered biomass production. This effect was likely caused by osmotic stress, as *K. petricola* growth is reduced upon addition of 1 M NaCl (Nai et al., 2013). The nitrogen content increased upon decreasing the supply of carbon or increasing the supply of nitrogen, whereas the carbon and hydrogen contents remained rather stable. The higher flexibility of the nitrogen content compared to the carbon content was also observed for other fungi, both when comparing species or conditions (Zhang and Elser, 2017; Camenzind et al., 2021; Pánek et al., 2024). At a medium carbon concentration of 273 mM and nitrogen concentration of 10.1 mM (i.e., a C:N ratio of 26.9), the model-inferred biomass C:N ratio of the WT was 17.7 ± 2.4 (95% CI), similar to the mean C:N ratio of 18 derived from a meta-analysis of 377 fungal species (Zhang and Elser, 2017). These ratios are lower for pathogenic fungi (mean C:N ratio of 13.04), followed by ectomycorrhizal fungi (C:N ratio of 14.44) and saprobic fungi (C:N ratio of 21.19) (Zhang and Elser, 2017). However, Pánek et al. (2024) observed generally lower C:N ratios for saprobes, soil saprobes having a distinctly lower C:N ratio (i.e., 7.0) than wood saprobes (i.e., 16.3). In general, the value for *K. petricola* therefore aligns best with ectomycorrhizal fungi and wood saprobes, indicating a natural environment characterized by nitrogen limitation.

Our measurements of the carbon and nitrogen content of the biomass did not distinguish intracellular from extracellular carbon and nitrogen (e.g., in EPS or cell wall components), nor whether these represented storage or non-storage compounds. If the former is the case, the increase of the nitrogen content under nitrogen surplus would be a type of surplus storage according to Chapin et al. (1990). Various carbon storage forms in fungi have been reported such as triacylglycerides, glycogen and trehalose (Elbein et al., 2003; Murphy, 2012; Mason-Jones et al., 2022). Nitrogen storage has been studied in filamentous fungi: the rapidly turning over cytosolic pool is dominated by glutamine and glutamate, while longer-term, high-density nitrogen reserves consist of basic amino acids (most notably arginine) frequently sequestered in vacuoles (Cramer et al., 1980; Kitamoto et al., 1988; Westenberg et al., 1989; Okreglak et al., 2023). Moreover, several fungi like *K. petricola* produce EPS, consisting of polysaccharides, proteins, nucleic acids and lipids, which may play various roles, including facilitating the attachment of colonies to substrata, buffering stresses, and modulating diffusion at the rock–air interface (Flemming and Wingender, 2010; Kirchhoff et al., 2017; Breitenbach et al., 2022). EPS have been proposed to function as an external storage pool, a view supported by the ability of *Candida* and *Botrytis* species to use their own EPS as a carbon source (Stahmann et al., 1992; Gientka et al., 2016). However, there is no evidence that RIF are able to re-use their EPS as a nutrient source.

The carbon source did not significantly influence biomass production, even though growth of other fungi was dependent on the carbon source being Glc or Suc (Itoo and Reshi, 2014). Depending on the species, fungi metabolize sucrose through extracellular or intracellular hydrolysis (Stambuk et al., 2000; Talbot, 2010; Marques et al., 2015). *S. cerevisiae* can hydrolyze Suc both extracellularly and intracellularly (Stambuk et al., 2000; Marques et al., 2015), whereas some plant pathogenic fungi take up Suc directly from the plant apoplast (Talbot, 2010). It is unknown which mechanism is utilized by *K. petricola*, but our observations show that *K. petricola* can hydrolyze sucrose without a noticeable metabolic cost.

The nitrogen source did, however, affect the biofilm morphology: 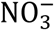 increased filamentation and penetration at intermediate and higher nitrogen concentrations, respectively. 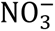-induced penetration was also observed in *Fusarium oxysporum* (López-Berges et al., 2010), *Fusarium graminearum* (Brauer et al., 2020) and *Magnaporthe oryzae* (López-Berges et al., 2010). Filamentous and ectomycorrhizal fungi prefer 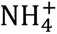 over 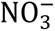 (with species-level variability) (Finlay et al., 1992; Keller, 1996). Because 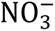 assimilation requires reduction, which diverts electrons from the respiratory chain and is thus energetically costly, 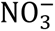 assimilation can lead to energy limitation. Hence, the enhanced filamentation and penetration under 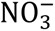 supply may represent a foraging response where fungi may extend hyphae more aggressively to explore the environment for 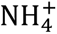 or other reduced nitrogen sources (Finlay et al., 1992; Hacskaylo et al., 2018).

Although melanization correlated with more filamentation, it did not affect biomass formation, indicating that melanin did not cause a growth deficit. (Catanzaro et al., 2024b) observed the same for *Cryomyces antarcticus*. Since biomass production by the WT was not reduced, we can reject our hypothesis that melanization increases sensitivity towards carbon limitation. The higher nitrogen content of the melanized WT (at higher nitrogen concentrations) is puzzling since the melanin of *K. petricola* is nitrogen-free DHN melanin, but may result from increased EPS production by non-melanized mutants shown previously (Breitenbach et al., 2022). Another possible explanation is a reduced chitin production by the mutant: a meta-analysis of 70 fungal species showed a positive correlation between nitrogen content and chitin content (Pánek et al., 2024). Chitin is essential for retaining/anchoring melanin in the fungal cell wall (Garcia-Rubio et al., 2020) and inhibition of DHN melanin synthesis in *Alternaria alternata* is accompanied by a reduced chitin synthesis (Fernandes et al., 2021).

Our observations have some important broader implications. The significant positive relationship between biomass and nitrogen availability suggests that these fungi could act as a pollution indicator as previously proposed by Krumbein and Gorbushina (2009). Furthermore, the increase of the fungal nitrogen content with increasing nitrogen supply suggests that subaerial biofilms mitigate nitrogen pollution in both agricultural (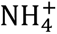) and urban (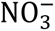) environments. A similar mitigating effect on nitrogen leaching has been attributed to soil fungi (de Vries et al., 2006).

In conclusion, we show here that carbon and nitrogen limitation affect the growth characteristics of *K. petricola* more than the carbon and nitrogen source and melanization. Surprisingly, our findings for the oligotrophic *K. petricola* generally agree with observations made for fast-growing fungi, suggesting that these characteristics are universal for fungi.

## Data availability statement

All data supporting the findings of this study are provided within the manuscript and the accompanying Supplementary Material. The R notebooks and their compiled HTML outputs have been deposited in Zenodo and can be accessed via the DOI: 10.5281/zenodo.18980984.

## Conflict of interest

The authors declare no conflicts of interest.

## Author contributions

**AD:** Conceptualization, Data curation, Formal analysis, Investigation, Methodology, Validation, Visualization, Writing – original draft, Writing – review & editing

**JS:** Conceptualization, Visualization, Writing – review & editing

**BJRC:** Formal analysis, Investigation, Methodology, Validation, Visualization, Writing – original draft, Writing – review & editing

**TC:** Conceptualization, methodology, data interpretation, review and editing

**KK:** Investigation

**JUK:** Conceptualization, Formal analysis, Methodology, Validation, Supervision, Writing – review & editing

**AAG:** Conceptualization, Visualization, Supervision, Writing – review and editing, Project administration.

**RG:** Conceptualization, Methodology, Visualization, Writing – original draft, Writing – review & editing, Supervision, Project administration, Funding acquisition

## Acknowledgements

We thank the Rillig Lab for giving us access to their LI-COR device. This work was funded by the Bundesanstalt für Materialforschung und -prüfung through the Ideas: Develop Program (Call 2021, IE2112).

## Notes

### Competing Interest Statement

The authors have declared no competing interest.

