## Supplementary Material for "Carbon and nitrogen availability affect biofilm growth and morphology of the extremotolerant fungus *Knufia petricola*"

**A**

| C Conc. (Glc) | **1 mM** | | | | |
|---|---|---|---|---|---|
| N Conc. ($NO_3^-$) | **N -** | **0.1 mM** | **1 mM** | **10 mM** | **100 mM** |
| C:N | — | **60** | **6** | **0.6** | **0.06** |
| Biofilm morphology | 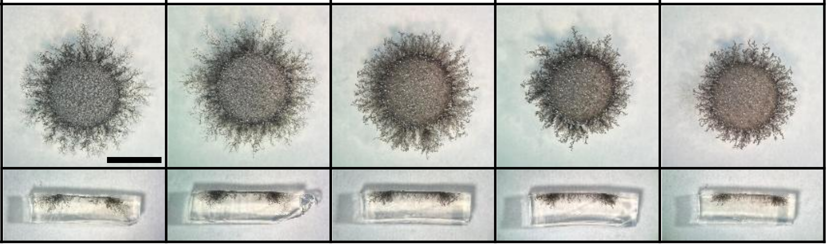 | | | | |

**B**

| C Conc. (Glc) | **C -** | **1 mM** | **10 mM** | **100 mM** | **1000 mM** |
|---|---|---|---|---|---|
| N Conc. ($NO_3^-$) | **0.1 mM** | | | | |
| C:N | — | **60** | **600** | **6000** | **60000** |
| Biofilm morphology | 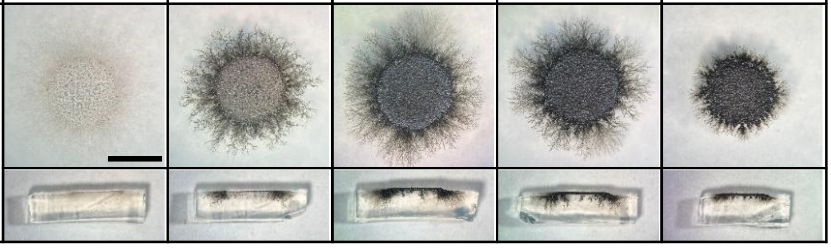 | | | | |

**Figure S1. Testing the co-limitation by both nutrients. (A)** Increasing the nitrogen concentration when carbon was limiting growth (based on the carbon experiment) did not affect biofilm growth or morphology. **(B)** Increasing the carbon concentration when nitrogen was limiting (based on the nitrogen experiment), in contrast, resulted in better growth at intermediate Glc concentrations and dual nitrogen and carbon limitation at the lower Glc concentration of 1 mM. In green are the conditions used in the carbon and nitrogen experiment (Figure 2). All experiments were conducted in triplicate. The scale bar represents 5 mm.

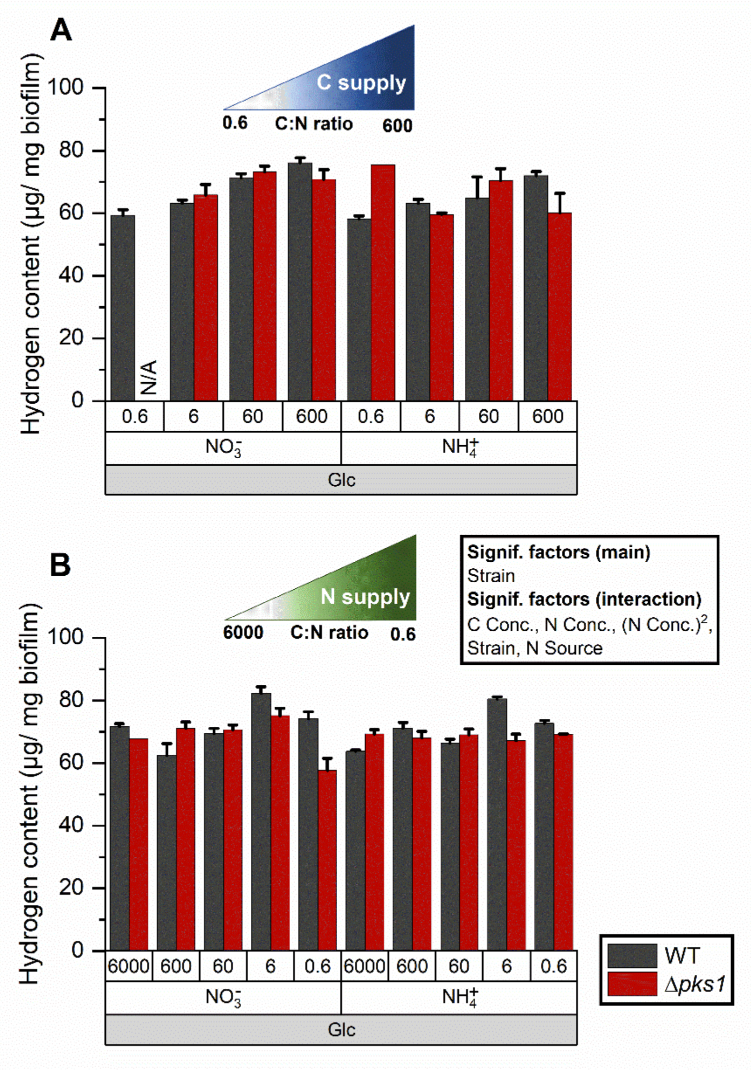


**Figure S2. Hydrogen content of the biomass for the (A) carbon and (B) nitrogen experiments after 28 days.** All experiments were conducted in triplicate; error bars show SE. Statistical significance is shown for the main variables and for the interactions among these variables based on the most parsimonious regression models (Table 2).

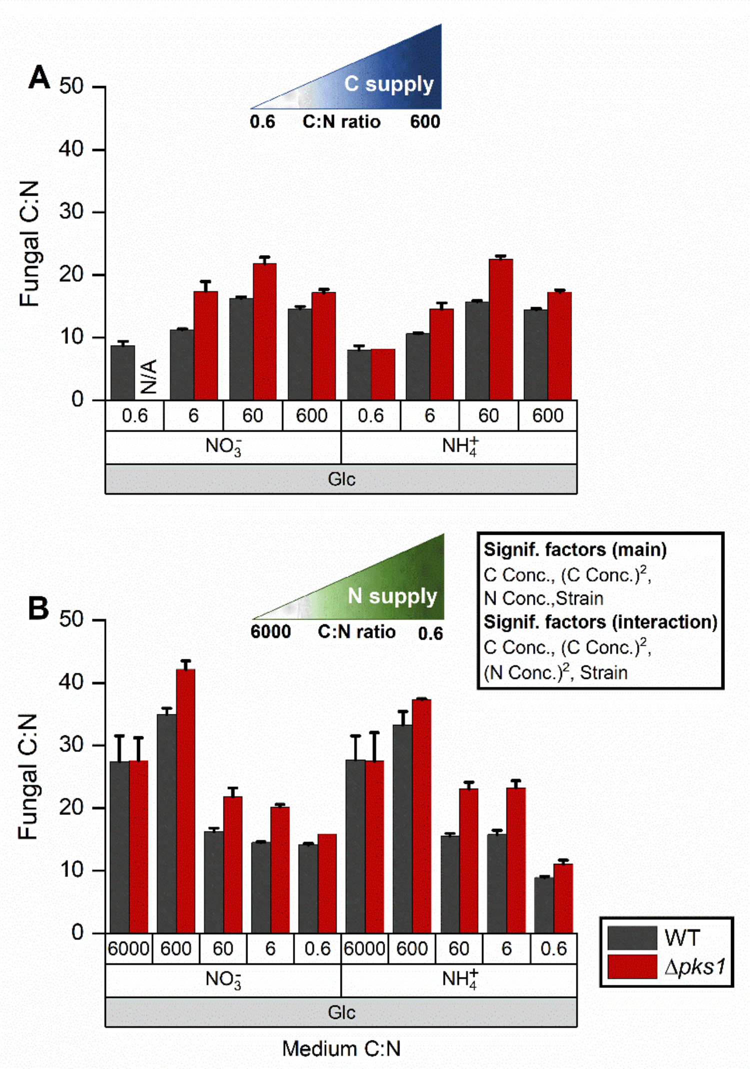


**Figure S3. Fungal biomass C:N ratio versus medium C:N ratio. (A)** With increasing carbon supply, the biomass C:N ratio increased to reach a maximum around the C:N ratio of 60. **(B)** For lower nitrogen supply, the biomass C:N ratio was much higher than in (A), dropping to ratios similar to (A) at higher nitrogen supply. All experiments were conducted in triplicate; error bars show SE. Statistical significance is shown for the main variables and for the interactions among these variables based on the most parsimonious regression models (Table 2).
